# Divergence in sleep architecture associated with interspecific shifts in memory investment

**DOI:** 10.64898/2026.11.07.765039

**Authors:** Euan Mortlock, Jessica Foley, Stephen H. Montgomery, Alice French

## Abstract

Sleep plays a critical role in stabilising memory. However, whether the architecture of sleep co-evolves with investment in learning and memory is unknown. To address this, we exploit the extreme levels of neural and behavioural divergence among Heliconiini butterflies. Compared to other Heliconiini, *Heliconius* have massively expanded mushroom bodies, the insect centres of learning and memory, which are associated with specific enhancements in visual learning and memory. Using a purpose-built video system, machine learning-facilitated behavioural tracking, and in-depth analysis of micro-behaviours during nighttime quiescence, we describe multi-state sleep in *Heliconius melpomene* and an outgroup Heliconiini, *Dryas iulia*. In both species, sleep is predominantly composed of “quiet sleep”, however, *Heliconius* and *Dryas* deviate in specific features. In *Dryas iulia,* quiet sleep is characterised by highly stereotyped, rhythmic antennal oscillations. Conversely, the antennae of *Heliconius melpomene* remain largely motionless but occasionally perform large movements which interrupt quiet sleep. These antennal deflections are associated with bouts of activity that include increased overall antennal displacement, isolated head twitching, proboscis movements and less frequently, wing movements. While these episodes were also seen in *Dryas iulia*, they were considerably less frequent. We further demonstrate that the behavioural profile of these “active sleep” bouts is constituently different from bouts of induced wakefulness, providing experimental support for multi-state sleep in Lepidoptera. This work shows that *Heliconius melpomene*, a species distinguished by its expanded mushroom bodies, exhibits more frequent bouts of active sleep than the outgroup species *Dryas iulia*. These findings raise the possibility that increased investment in learning and memory, as well as the structures that support it, may be associated with changes in sleep architecture.

## Results

*Heliconius melpomene* and *Dryas iulia* were selected as representative species of the divergence in neural and behavioural traits across the radiation of Heliconiini butterflies. Compared to outgroup Heliconiini, such as *D. iulia*, *Heliconius* butterflies, including *H. melpomene*, have massively expanded learning and memory centres (mushroom bodies) with 8X more intrinsic neurons (Kenyon cells)^1^, associated with elongated lifespans^2^ and an improved capacity for long-term visual memory and combinatorial learning^1,3,4^. These traits are associated with a novel diet to exploit protein- and fat-rich pollen grains, and a linked foraging strategy more dependent on long term spatial memory^5,6^. *H. melpomene* and *D. iulia* represent the extremes of this variation, providing a framework for in-depth exploration of their sleep. Within this framework, we therefore sought to describe nighttime sleep in Heliconiini and asked whether antennal movements have a unique behavioural signature. Secondly, we asked whether there is evidence for different sleep states, and if these vary between species with high vs. low neural investment in learning and memory.

### Nighttime quiescence and posture changes characterise Heliconiini sleep

Using video tracking to capture 24-hour rest/activity patterns in *D. iulia* and *H. melpomene* (Figure 1A), we found that both species show clear diurnal activity patterns (Figure 1B,C). In both species more activity was observed during the light phase compared to dark (Wilcoxon rank sum test, p < 0.05 for both *H. melpomene* and *D. iulia*). Even though both species were mostly inactive during the dark phase, *H. melpomene* still displayed more dark-phase movements than *D. iulia* (Wilcoxon rank sum test, p < 0.05, Figure 1C). From these recordings we identified two hallmarks of sleep. First, each individual remained largely sedentary for the duration of the dark phase^7^. Second, in most animals, sleep is associated with postural change, which in insects can involve relaxation of neck tone, drooping of the head, and lowering of the body towards the substrate, no longer resisting gravity ^8–10^. Both species exhibited postural relaxation at the onset of nighttime quiescence (Figure 1D) and, when sleeping hanging upside down, were further from the ceiling substrate at ZT23 compared to ZT12 (Directional Wilcoxon signed rank test, p < 0.05, Figure 1E).

**Figure 1:**
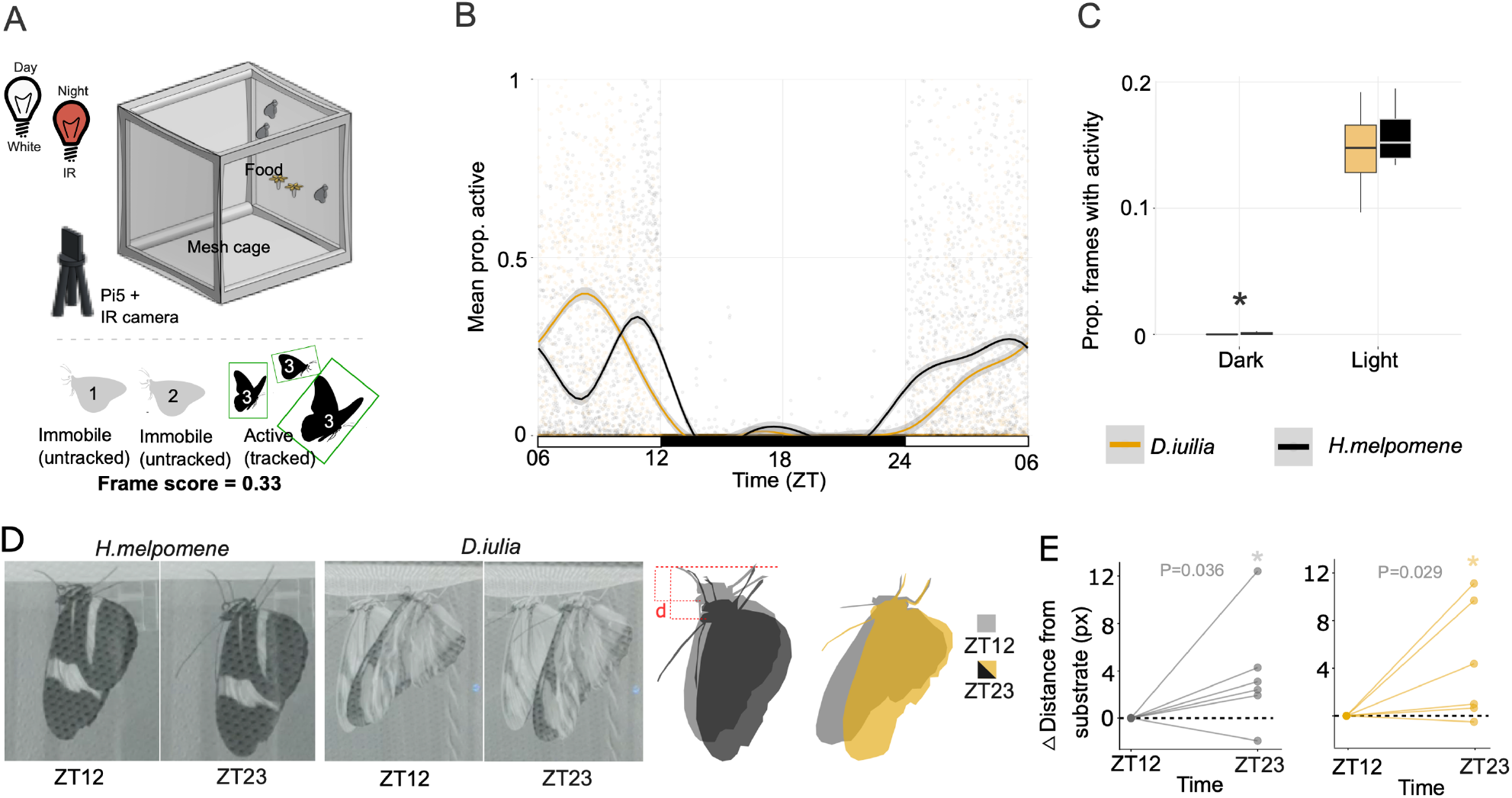
*D. iulia* and *H. melpomene* are diurnal species showing consolidated periods of inactivity at night. A) Top: schematic of the cage setup for 24-hour activity monitoring. Bottom: Calculation of frame-level activity score based on pixel-luminosity frame differencing. B) Activity profile showing mean number of butterflies active per minute over 24 hours in *D. iulia* (yellow) and *H. melpomene* (black). Lines were smoothed using GAM and x-axis shows zeitgeber time (ZT). C) Proportion of frames detecting activity in *D. iulia* (yellow) and *H. melpomene* (Black) during the dark and light phases. D) Example images and outlines of *H. melpomene* (Left) and *D. iulia* (Right) at ZT12 and ZT23. E) Quantification of change in vertical distance from centre of eye to where front tarsus contacts mesh substrate from ZT12 to ZT23. N=6, one-sided Wilcoxon Rank-sum (*P<0.05).

### Pronounced micromovements during Dryas iulia sleep

Detailed descriptions of insect micromovements during sleep have previously shown that antennae are typically relaxed and inactive relative to wake^9,11,12^. However intermittent quivers^13^, rhythmic movement^12^, and bursts of activity^14^ have been reported in a number of species. Therefore, we next explored the presence of sleep-specific antennal micromovements in *D. iulia* and *H. melpomene.* Using bespoke behavioural chambers (‘*Papilyawns’*; Figure 2A), high resolution video and automated tracking^15^ of antennal movements, we captured antennal coordinates throughout nighttime sleep (Figure 2B). From these coordinates, we used spectral analysis (STAR methods) to describe the nature of antennal movements during 20 second windows and observed that the dark-phase antennal movements of *D. iulia* are characterised by small rhythmic oscillations (Figure 2B; Figure 2D). These oscillations have a frequency 0.2–1.5 Hz with a dominant frequency of 0.5Hz (Figure 2E, Supplementary Video 1), with spectral concentrations and band power values being higher in the dark phase than in light (W-D/W-L = 0.89 vs 0.32, power = 79.6 vs 0.03 px² Hz⁻¹, Figure 2F). These observations were evident at the population level, with median ACF at lag (STAR methods) being significantly higher during the dark phase compared to the light (Wilcoxon signed rank, P < 0.01, Figure 2G), meaning antennal oscillations were dark phase-specific in *D. iulia* (Figure 2H). In *H. melpomene,* windows meeting oscillation criteria (STAR methods) were, in contrast to *D. iulia*, rare, fragmented and not strictly restricted to the dark phase (Figure 2H). Instead, antennae spent most of the time largely immobile (Supplementary Video 2), interrupted by large deflections (Figure 2B-D). At the population level there was no significant difference in median total displacement of antennae per window between dark and light phases in *H. melpomene* (Wilcoxon signed rank, P = 0.677). However, median variance per window did differ significantly (Pitman-Morgan, P = 0.044) indicating the nature of movements during sleep were different from wake in *H. melpomene* (Figure 2I).

**Figure 2:**
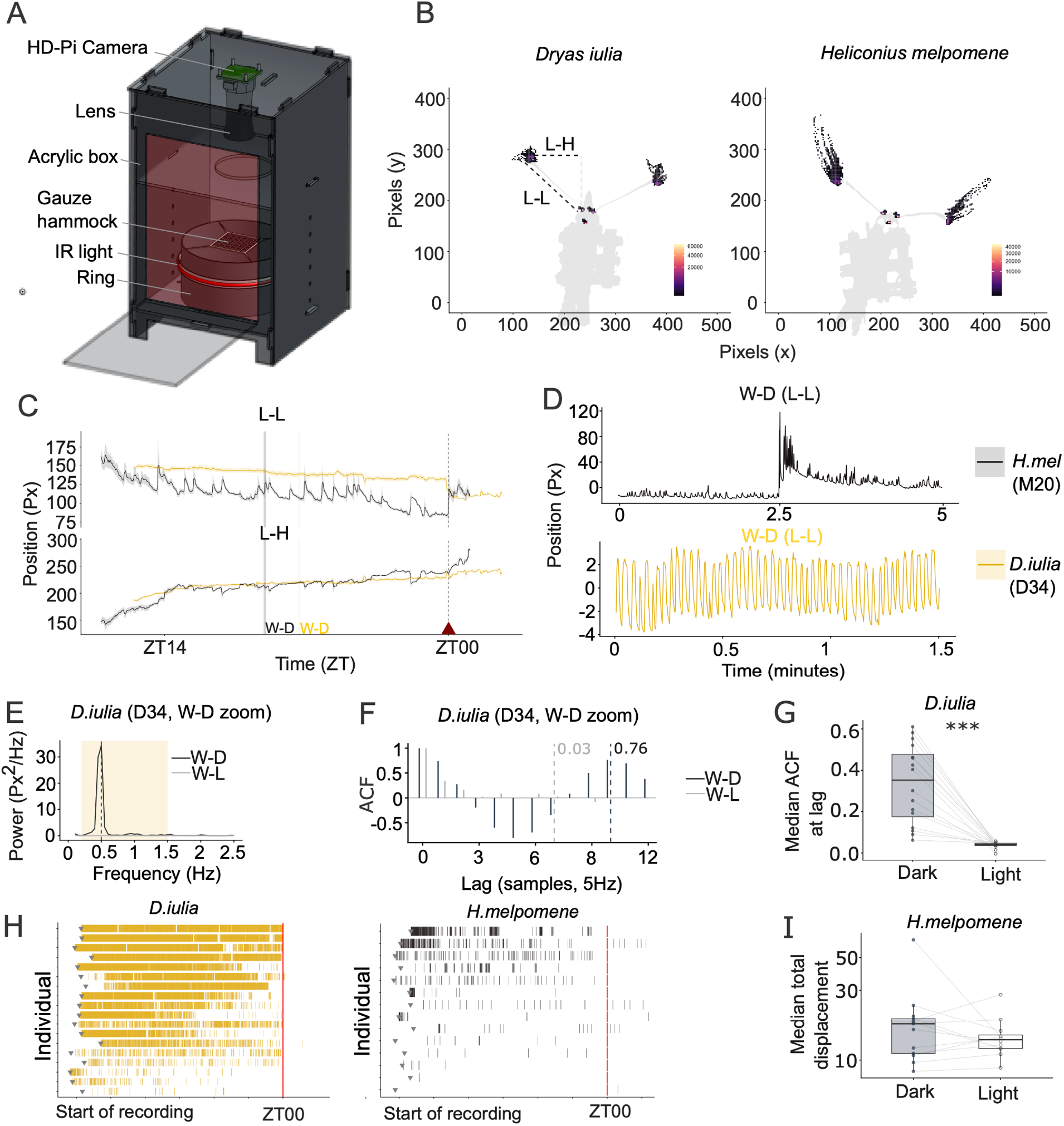
**D. iulia and H. melpomene show species-specific antennal movements during nighttime sleep.** A) CAD of ‘Papilyawn’ sleep chambers. B) DLC tracked coordinates of antennal bases, clubs and proboscis for a representative *D. iulia* (left) and *H. melpomene* (right) between zeitgeber time (ZT) 18-19. Coordinates derived from DLC video annotation. C) Traces show 5-minute rolling means for L-L (top) and L-H (bottom) channels from representative individuals for *D. iulia* (yellow) and *H. melpomene* (black). Ribbons show +/- median absolute deviation. Grey/yellow vertical lines mark W-D. D) Expanded W-D for *H. melpomene* (Top) and *D. iulia* (bottom). Signals are locally mean-centred for each window, dark-phase epochs selected to illustrate oscillation coherence. E) Power spectrum density (PSD) of W-D and W-L; shading = 0.2-1.5Hz band. F) Autocorrelogram of detrended signal in dark and light phases (W-D and W-L). Dashed lines mark the lag of one oscillation cycle, annotated with autocorrelation coefficient at a lag of one oscillation cycle (ACF). G) Per-individual median ACF at lag in the dark and light phases, *D. iulia* (N = 16); lines join individuals. Paired Wilcoxon signed-rank test (*** p < 0.001). H) Sustained oscillation rasters for all recorded individuals: *D. iulia* (left, N = 18) and *H. melpomene* (right, N = 15). One individual per row; each bar represents one 20-second sliding window epoch that passes the matched filter (spec_conc ≥ 0.55, acf_at_lag ≥ 0.10). I) Per-individual median antennal displacement in light and dark phases, *H. melpomene* (N = 14). Paired Wilcoxon signed-rank test for medians; Pitman-Morgan test for variances (*** p < 0.001, ns P > 0.05).

### Multi-state sleep and increased frequencies of active sleep in Heliconius melpomene

Variation in the presence and frequency of antennal micromovements has been interpreted as evidence for different sleep states in arthropods. Further, attempts to correlate antennal movements with electrophysiological recordings has revealed that during “quiet” sleep neural activity is low, and during “active” sleep it is typically higher^12,16^. To explore sleep states in *Heliconius* and *Dryas*, we looked for inconsistencies in full night antennograms, which we defined as changes to baseline antennal behaviour, for example large antennal deflections in *H. melpomene*. We then manually scored frames proceeding these events, recording the timing, frequency and duration of active episodes (Figure 3A)^17^. Overall, the total number of active sleep bouts per individual was significantly higher for *H. melpomene* compared to *D. iulia* (Figure 3B; Wilcoxon rank-sum test, P = 0.001). Next, we quantified behaviour during these bouts, comparing antennal movements pre, during and post a putative active sleep bout (Figure 3C). Active bouts included incidences of proboscis, head, and body movements, the majority of which could be described as ‘twitching’ (Supplementary Videos 3 and 4). Less frequently we observed wing opening and flapping. While it was not possible to apply power spectrum density (PSD) analysis to active sleep bouts due to their short duration, we did quantify total antennal displacement and observed that in both *D. iulia* and *H. melpomene*, active sleep bouts were associated with more movement relative to the same duration of time preceding it (Figure 3D, Wilcoxon signed rank test, *D. iulia*: P = 0.049 and *H. melpomene* P < 0.001).

**Figure 3:**
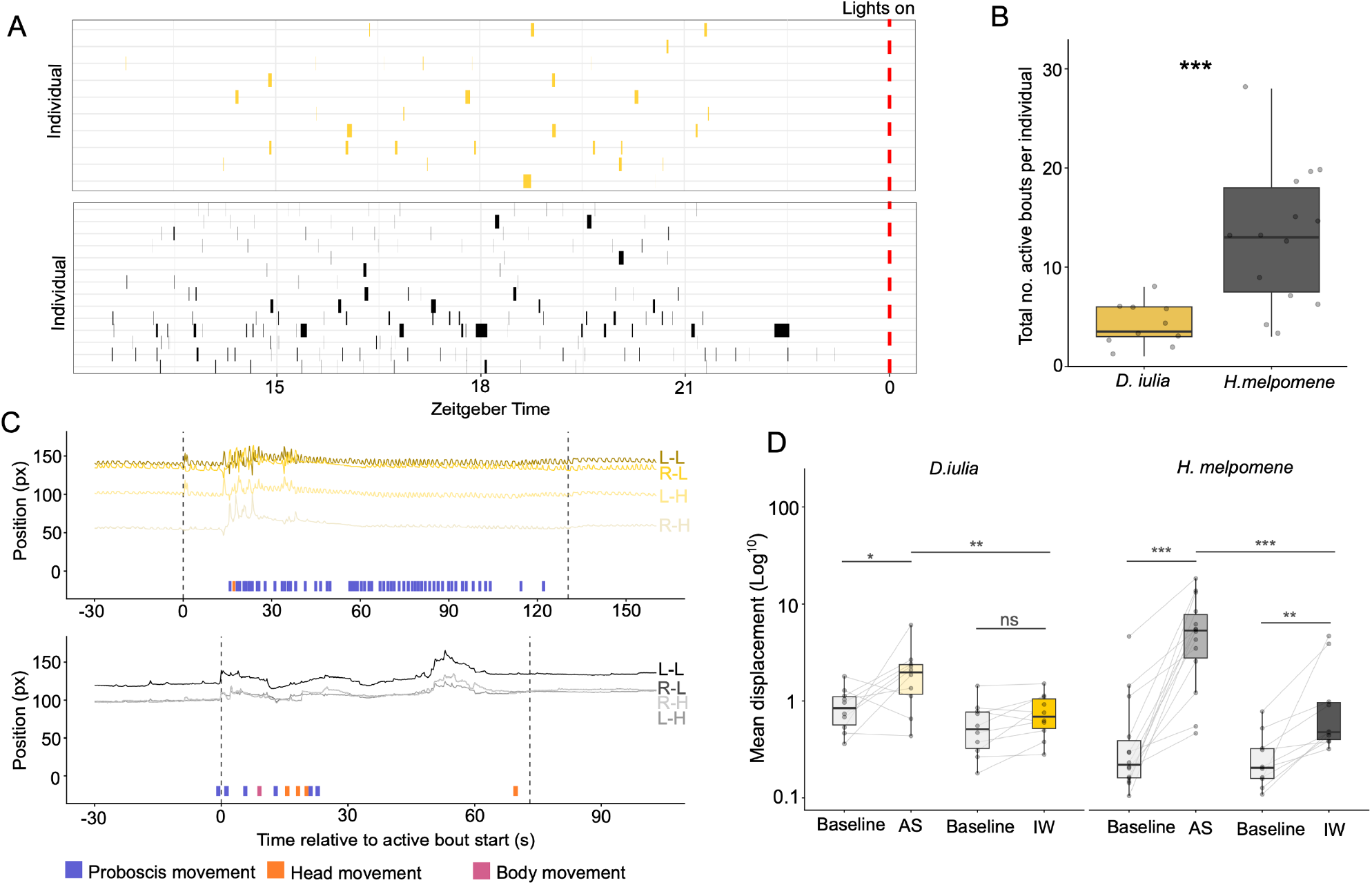
**H. melpomene show more active sleep bouts than D. iulia.** A) Timing and duration of active sleep bouts across the night for all *D. iulia* (top) and *H. melpomene* (bottom); one row per individual; red dashed line, lights on. B) Total active sleep bouts per individual in *D. iulia* (yellow) and *H. melpomene* (dark grey). N=10-14, Wilcoxon rank-sum test (*** p < 0.001). C) Antennal channels around an active sleep bout in a representative *D. iulia* (top) and *H. melpomene* (bottom). Dashed lines mark bout start and end times; ticks mark scored proboscis (blue), head (orange) and body (pink) movements. D) Mean antennal displacement (best channel, log scale) during active sleep (AS) and induced waking (IW) with matched baseline in *D. iulia* (Left) and *H. melpomene* (Right). Lines join individuals (Paired Wilcox test, * p < 0.05, ** p < 0.01, *** p < 0.001, ns p > 0.05).

To strengthen our resolve that active bouts are a distinct sleep state rather than spontaneous awakenings, we performed a final set of experiments whereby sleeping butterflies were woken with a mechanical stimulus delivered by a vibrating disc (Supplementary Videos 5 and 6). Following the stimuli, we quantified antennal movements and the frequency of other behaviours observed during active sleep bouts. Initially, mechanical stimulation induced an interruption to baseline antennal activity (Supplementary Figure 2) and increased mean antennal displacement in *H. melpomene* (Wilcoxon signed rank, P = 0.002) but not in *D. iulia* (Wilcoxon signed-rank, P = 0.064) during the post-stimulus period (Figure 3D). In both species, mean antennal displacement was significantly lower compared to the displacement seen during active sleep (Wilcoxon signed rank *H. melpomene* P=0.001; *D.iulia* P=0.009). The behavioural profile of active sleep bouts was also distinct from post-vibration induced waking (Figure 4A,B). In both *D. iulia* and *H. melpomene,* compared to induced waking, active sleep was associated with significantly more isolated head movements (Wilcox test AS vs IW *D. iulia* P = 0.001; AS vs IW *H. melpomene*, P = 0.002) and proboscis movements (Wilcox test AS vs IW *D. iulia* P= 0.005; AS vs IW *H. melpomene*, P=0.002). Frequencies of wing flapping were similar between active sleep and induced waking (Wilcox test AS vs IW *D. iulia* P= 0.49; AS vs IW *H. melpomene*, P=0.74), whereas wing opening was more common following induced waking compared to active sleep in *H. melpomene* (Wilcox test AS vs IW *D. iulia* P = 0.051; AS vs IW *H. melpomene*, P = 0.001). Arousal following mechanical stimuli also nearly always involved postural adjustment (rapid restoration of body tone, returning towards the substrate; Supplementary Videos 5 and 6), whereas active sleep bouts did not involve postural change. Together the distinct behavioural profiles presented here suggest that spontaneous active sleep bouts are constituently different from induced waking and are more frequent in *H. melpomene* than *D. iulia*.

**Figure 4:**
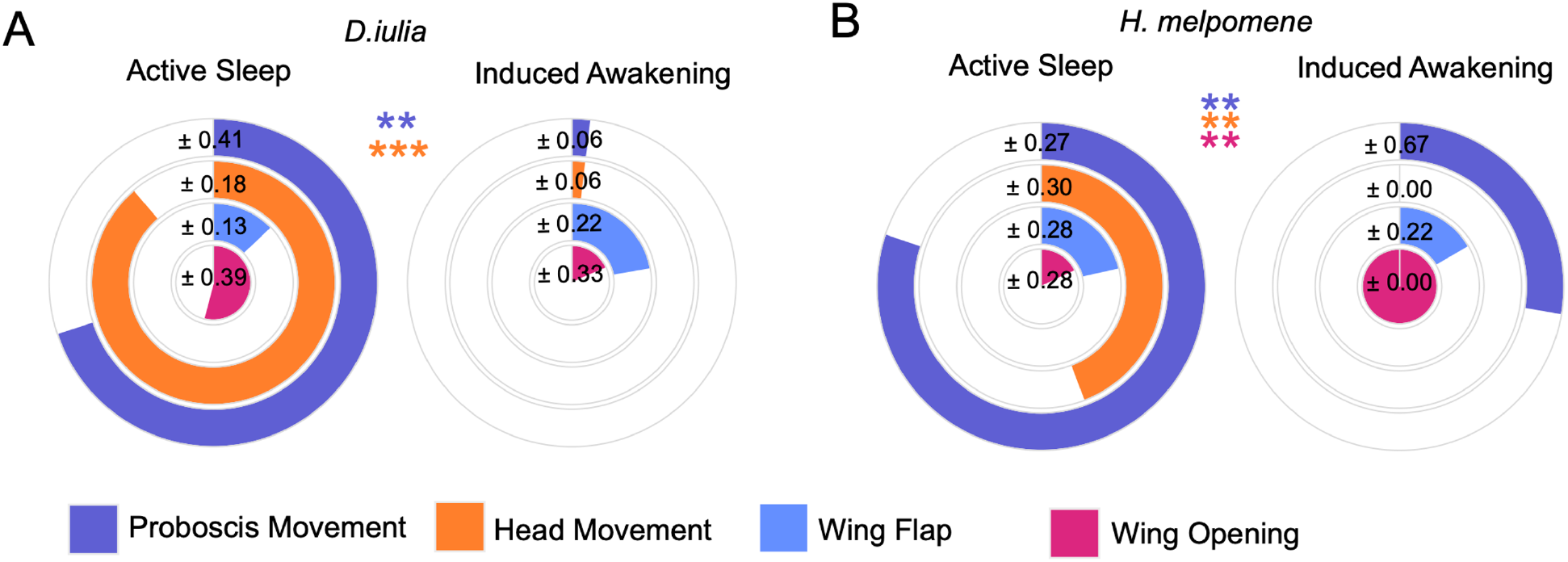
**Behavioural profile of butterflies in active sleep is distinct from wake.** Concentric ring charts show the mean proportion of active sleep bouts (left) and induced waking periods (right) containing at least one instance of proboscis movement (purple, outer ring), head movement (orange, 2^nd^ ring), wing flapping (blue, 3^rd^ ring), and wing opening (pink, inner ring) in *D. iulia* (A, left; n = 10 active sleep, 9 induced waking) and *H. melpomene* (B, right; n = 14 and 9). Values are ± SD across individuals. Wilcoxon rank-sum test used to compare behavioural profiles between states (**P<0.01, *** p<0.001).

## Discussion

In recent years, efforts to characterise sleep in a diverse range of species^8,18–21^ have not only revealed its ubiquity across the animal kingdom but also how its form and function have evolved across short evolutionary timescales. Here, we provide a novel behavioural characterisation of nighttime sleep in untethered Heliconiini butterflies, *D. iulia* and *H. melpomene*, which have pronounced divergence in adult life history and investment in learning and memory^1,4,5^. We show that these diurnal species remain largely quiescent throughout the night and display postural relaxation, two established hallmarks of sleep^7^. Through analysis of micro behaviours, we provide evidence for two distinct sleep states that are distinguishable from wakefulness and one another.

During most of the night, the antennae of *D. iulia* are rhythmic, oscillating consistently, a behaviour that, as far as we are aware, has not yet been described in other Lepidoptera. In *Aglais* butterflies and pyralid moths, during rest antennae remain motionless and folded back under the wings^22,23^. Conversely, *D. iulia* quiet sleep-antennal oscillations cease during wakefulness and during active sleep bouts. Active sleep bouts are characterised by a deviation from quiet sleep baseline antennal movements (in *D. iulia*, oscillations cease and antennal movement becomes irregular; in *H. melpomene*, otherwise motionless antennae make large, non-rhythmic deflections), head twitches, proboscis pumping and on occasion wing opening/flapping. Proboscis movement^12,24^ and a related behaviour, ventilatory bursting^9^ have been observed during sleep in bees, flies and now butterflies. It is proposed these behaviours drive haemolymph flow, necessary to combat poor circulation after long periods of immobility and to clear metabolic waste, a process analogous to glymphatic clearance in mammals^24,25^. Head twitching was also frequently observed during active sleep, which could be related to replay in sensorimotor circuits^26^.

A second notable finding was that in *H. melpomene*, active sleep bouts are more prevalent than in *D. iulia* and punctuate otherwise largely motionless quiet sleep. Mammalian phylogenetic analyses addressing the relationship between increased relative brain size and sleep composition provide mixed evidence^27–29^ Nonetheless, we propose the shift in multi-state sleep between *H. melpomene* and *D. iulia* is related to increased investment in the mushroom bodies, the insect learning and memory centres. *Heliconius* mushroom bodies contain eight times more neurons than *Dryas*^1^, supporting specific shifts in learning and memory^1,3,4^, and visual long term memory stability that remains robust across their elongated lifespans^30^. Importantly, despite this suite of derived traits, *Heliconius* and *Dryas* are closely related (Supplementary Figure 1), share common broader ecologies, juvenile life histories, hostplants and habitat preferences. Any differences between these species are therefore most plausibly related to the unique diet, life history and neural investment of *Heliconius*. However, as these traits arose together in the *Heliconius* lineage, a comparison of two species cannot definitively attribute differences in sleep architecture to any one of them; broader sampling across Heliconiini, where mushroom body size and foraging ecology vary among species^31^ will help delineate this.

The mushroom bodies regulate sleep^32^ and support memory formation. Following release from a novel site, honeybees sleep more upon return to the hive and the retention of learned homing routes is hampered if sleep is disrupted ^32^. *Drosophila* also sleep more following courtship learning tasks ^34^ and social enrichment ^35^ indicating that wakeful experience may influence sleep need. Innate scototactic behaviours in cockroaches can be overridden through associative shock conditioning, consolidation of which is enhanced if “rest” follows^36,37^. The mechanistic explanation for how sleep supports memory consolidation is well established; in *Drosophila* ^18,37^, zebrafish ^39^ and mice ^40^ sleep selectively preserves or downscales synaptic boutons, potentially following a “use-it or lose-it” basis, a process that is impaired following deprivation.

Given that synaptic remodelling has functional implications for memory consolidation^41^ and is proposed as a key function of sleep, animals with high cognitive capacity and corresponding neuronal enhancement should increase investment in sleep, either by increasing total sleep duration or by altering its composition (such as more time in a particular sleep state)^42^. Our data are consistent with this hypothesis, suggesting increased frequency and prevalence of active sleep may have co-evolved with mushroom body expansion in Heliconiini butterflies. This raises the tantalising possibility that active sleep, which may carry out functions distinct from quiet sleep, has expanded to support memory related processes in *Heliconius*. Whether this is a generalisable feature of mushroom body evolution, and whether sleep composition responds to daily cognitive demand in *Heliconius*, is of clear future interest.

## Resource availability

Requests for further information should be directed to and will be fulfilled by the lead contacts Stephen Montgomery and Alice French.

## Materials availability

This study did not generate any new, unique reagents.

## Data and code availability

All data and code will be available at: 10.5281/zenodo.23541322

## Acknowledgements

We are grateful for the support of a Research Grant from the Association for the Study of Animal Behaviour (ASAB) to SHM/AF, and an EPSRC grant (Horizon Guarantee Scheme-EP/Y032071/1) to AF.

## Author contributions

SHM and AF designed the study and acquired funding. All authors contributed to the study execution, data analysis and writing.

## Methods

### Experimental model and study participant details

*Heliconius melpomene* and *Dryas iulia* pupae were obtained from commercial suppliers (The Entomologist Ltd) and pupae derived from stock populations established from the same supplier. Pupae were incubated in climate-controlled rooms at 25°C and 80% humidity under 12:12 light:dark conditions. Upon eclosion, butterflies were transferred to 2m x 1m x 1m (L,H,D) mesh cages in the same room. Adult butterflies were fed a solution containing 20% v/v sugar, 5% v/v critical care formula (Vetark, CCF15) from artificial feeders. The solution was refreshed every 2 days. As a pollen source, butterflies were also provided with fresh *Lantana camara* flowers which were changed every 4 days.

### 24- hour activity monitoring in mesh cages

To capture *D. iulia* and *H. melpomene* daily rest/activity rhythms in freely moving individuals, butterflies were placed either singly or in groups (n = 2-3) into mesh cages (30cm x 30 cm x 30 cm) 2-4 hours before onset of the dark phase (n = 4 cages total per species). Cages were kept inside climate-controlled rearing chambers to maintain light, temperature and humidity conditions. Infrared lighting allowed tracking through the dark phase, and two feeding stations allowed butterflies to feed *ad libitum.* An infrared camera (NoIR v3) connected to a Raspberry Pi 5 was used to capture a 24- hour video of the cage starting from when butterflies were introduced (Figure 1A). Resolution and frame rate were 640 x 320 and 30fps. Scripts used for video capture and instructions for mounting a hard drive are included in the data repository.

### Sleep recordings in Papilyawns

Sleep recordings were obtained from both male and female individuals aged between 6 and 24 days post eclosion with an HD Raspberry Pi camera under infrared illumination. Butterflies were collected from rearing chambers approximately 1 hour after lights off, when quiescent, and were encouraged to hang on the underside of a gauze hammock inside the recording chambers (‘Papilyawns’; Figure 2A). The specifications and cost of materials for construction of Papilyawns are detailed in the Data Repository. Once settled, recordings were initiated and were stopped after lights on (approximately 13 hours later).

### Inducing waking with vibration

In a separate set of nighttime video recordings using Papilyawns, hammocks were fitted with 2 vibrating disks to deliver gentle and strong vibrations to butterflies. 6 vibrations were delivered for 3 seconds to butterflies at 105 minutes intervals, the first timed 105 minutes from the onset of recording. The order of vibrations was [strong, gentle], [gentle, strong], [strong, gentle]. We observed no difference in response other than inducement of waking to strong/gentle vibrations, so all were pooled in analyses. Raspberry Pi Picos were used to time the stimuli: bill of materials, code and instructions for Pico configuration are included in the Data Repository. Behavioural responses following vibrations were scored using BORIS software for the 60s following vibration and exported as CSV files.

### Video analysis of free flying butterflies

24-hour videos were split into three sections (light phase 1, dark phase and light phase 2) with Shotcut (Meltytech LLC) and subsequently processed using the python library of OpenCV (OpenCV team). Initially, videos were converted to greyscale, and each pixel was assigned a luminance value from 0-255 (black-white). A mask was applied to the video and pixels within the region of interest (ROI) were binarized using a threshold of 127. To generate a cage-level readout of movement (did any one animal move) we used a frame differencing approach: pixels where luminosity varied more or less than 15 (proxy for movement, set to exceed threshold for noise), relative to the preceding frame, were dilated with a 3x3 grid to encompass neighbouring pixels also meeting this threshold, generating clusters. Because the classifier initially struggled with butterflies rapidly changing shape and size during flight, we increased pixel connectivity to allow diagonal as well as orthogonal dilation. Clusters were then filtered to eliminate those smaller than 100px^2^ and greater than 4000px^2^ (encompassing smallest and largest butterfly surface area). Binary images showing threshold-met pixel luminosity changes were generated for each frame. Frames containing cluster(s) were assigned 1 (movement detected) otherwise 0 (no movement). This information, along with frame number, time (seconds from start of video), number of clusters, their surface area and total cluster surface area were compiled into a master .csv file and subsequently analyzed in R (v. 4.6).

Initially tracking data was stitched and aligned to zeitgeber time (ZT). Raw data at 30 rows per second was under sampled to 1 row per second by taking the maximum activity score per second. We then calculated the proportion of frames (within 1-minute bins) containing activity (prop_active) and normalized according to number of butterflies present in the cage (prop_active_n, Figure 1B). While our overall detection of movement at the cage level was robust, in some frames, typically when in flight, we observed that a single butterfly can generate multiple clusters, making our estimation of the *number* of butterflies active, less reliable. However, we reasoned that frame averaging would sufficiently dilute these instances to give a reasonable readout of number of individuals active.

### Postural analysis

At dusk (∼ZT11-12), already quiescent butterflies were transferred from communal colonies to smaller terrarium where they were encouraged to hang upside down from the custom-designed mesh shelf and were recorded overnight. The majority remained *in situ* but those significantly shifting positions were excluded from analysis (N=2, *Dryas iulia*). Snapshots were extracted from videos 1 hour following onset of recording (ZT12-13) and again before lights on (ZT23). Using ImageJ, the vertical distance from the centre of the eye to in line with where the front tarsi contacted the mesh ceiling/substrate was used to quantify postural relaxation in each extracted frame.

### Video analysis using DeepLabCut (DLC)

To gain an understanding of sleep-specific microbehaviours in *H. melpomene* and *D. iulia*, 63x ∼13 hour overnight videos were recorded capturing both undisturbed sleep and induced waking experiments for individual butterflies. Videos were recorded in 480p at 30 frames per second. Frames from these videos were then annotated and used to train the DLC pose estimation model^15^, which was then run on all videos to track antennal movements during periods of sleep. In brief, 10x frames were extracted from each of the 63 videos using the default k-means clustering method, and the DLC graphical user interface was then used to manually label 5 body parts for each of the 630 frames: left club (the tip of the left antenna; clubL), right club (clubR), base of left antennae (baseL), base of right antennae (baseR), and the mouthparts (proboscis). A skeleton was then applied linking baseL-clubL and baseR-clubR to define each antenna as a “bone”. The DLC model was then trained on these labelled frames with a single shuffle, using the default settings with the following changes: crop_sampling = FALSE, motion_blur = TRUE, batch_size = 8, epochs = 200. hflip was also set to TRUE, with p: 0.5 to apply a horizontal flip with 50% probability, and clubL/clubR and baseL/baseR defined as symmetric body parts. The trained model’s performance was then evaluated by visually comparing manual labels for each body part and the predicted labels applied by the model, and ensuring the average root mean square error between these labels was <5 pixels. Once deemed satisfactory, this trained model was then used to analyse all 63x 13-hour videos in full, producing annotated videos and .csv files containing the xy coordinates of all labelled body parts as well as the antennal “lengths” (as a proxy for z antennal movement), over time. These computationally-expensive model training and video analysis steps were conducted using the BlueCrystal high-performance computing cluster at the Advanced Computing Research Centre at the University of Bristol. The resulting .csv files were filtered using the deeplabcut.filterpredictions command run on defaults and analysed in R (v. 4.6) to assess species-specific nature of antennal movements during undisturbed nighttime inactivity, or pre/proceeding a disturbance (vibration).

### Quantification of DLC data

To identify and characterise any consistent patterns of movement in antennae, we applied fast Fourier transform (FFT) to raw data. Filtered DLC coordinates were converted to four antennal channels per frame; antennal length (base to club distance, L-L/R-L) served as a z-dimension proxy; horizontal base to club distance (L-H/R-H) captured lateral movement isolated from any small body movements. Raw position data was downsampled from 30 to 5Hz by taking every 6^th^ frame. Data were binned to 20 second windows (100 samples) with a 10 second slide and linearly detrended to further dampen any body movements. For each window we derived the dominant frequency (>0.1Hz), power spectrum density (PSD, Welch’s method), band power (sum PSD within 0.2-1.5 px² Hz⁻¹) and spectral concentration (proportion of power above 0.1 Hz within px² Hz⁻¹) and spectral concentration (proportion of power above 0.1 Hz lying within ±0.2 Hz of the dominant frequency; ∼1 for a pure oscillator and ∼0 for noise). We scored the rhythmicity of oscillations as the autocorrelation coefficient at a lag of one oscillation cycle (ACF at lag 1). Windows whose dominant frequency fell below the 0.1-Hz floor were excluded, because the lag equalled half the window and ACF at lag became unreliable. For each individual, a single antenna channel was used for analysis (for visualisation multiple channels are used), where each channel was scored (for every window, median spectral concentration*median amplitude interquartile range) and the channel with the highest score across all windows was selected, per individual. Scoring channels in this manner quantifiably identified the most focused and largest oscillations, and matched visually labelled oscillations in both *D. iulia* and *H. melpomene*. We classified a window as oscillating when spectral concentration and ACF at lag exceeded 0.55 and 0.10 respectively, with thresholds derived from visually labelled oscillatory bouts in both species. To restrict detection to sustained behaviour, only runs of ≥ 3 consecutive passing windows were retained (Figure 2H). Windows after lights on were not suppressed, thus sparse post-lights-on detections in *D. iulia* reflect cessation of oscillation on waking. Data analysed in the dark phase ran from 30 min after recording start to 30 min before lights on; the light phase ran from lights on to the end of recording. Per-individual medians were calculated for each phase, and two *D. iulia* were excluded from this comparison due to flying out of camera before lights on (Figure 2G).

### Quantification of active sleep

Putative active sleep bouts were identified in full night antennograms (included in the Data Repository) as stereotyped departures from an individual’s baseline activity, as large deflections (*H. melpomene)* or cessation of oscillation (*D. iulia*). In BORIS, each video was reviewed at 8x speed to confirm deflections represented in antennograms were legitimate events, and detect any events not present in antennograms. Event onset was scored at the first frame of departure from baseline and offset once active behaviours had ceased for >10s. Point behaviours of proboscis, head, body and wing movement were scored within bouts. All timestamped active events were exported from BORIS as .csv files and used for subsequent analysis in R. Paired Wilcoxon signed-rank tests were used to compare medians; Pitman-Morgan tests were used to compare variances.

### Data repository contents

| **REAGENT or RESOURCE** | **SOURCE** | **IDENTIFIER** |
|---|---|---|
| **Deposited Data** |  |  |
| Video | Recorded in the lab | Supplementary_video_1_Dryas_iulia_D36_QS.mp4 |
| Video | Recorded in the lab | Supplementary_video_2_Heliconius_melpomene_M22_QS.mp4 |
| Video | Recorded in the lab | Supplementary_video_3_Dryas_iulia_D36_ASB.mp4 |
| Video | Recorded in the lab | Supplementary_video_4_Heliconius_melpomene_M22_ASB.mp4 |
| Video | Recorded in the lab | Supplementary_video_5_Dryas_iulia_D50_IW.mp4 |
| Video | Recorded in the lab | Supplementary_video_6_Heliconius_melpomene_MR8_IW.mp4 |
| STL (3D print) | Designed by EM | Frame_connector_part_1_Figure_1.stl |
| STL (3D print) | Designed by EM | Frame_connector_part_2_Figure_1.stl |
| DXF (Laser cut) | Designed by EM | Base_laser_cut_Figure_1.dxf |
| DXF (Laser cut) | Designed by EM | Frame_laser_cut_Figure_1.dxf |
| STL (3D print) | Designed by AF | Terrarium_shelf_part_1_Figure_1 |
| STL (3D print) | Designed by AF | Terrarium_shelf_part_2_Figure_1 |
| Bill of Materials for Figure 1 (.xlsx) | Source of each product specified in the spreadsheet | Bill_of_materials_Figure_1.xlsx |
| R script | Written by AF | Figure_1E.R |
| Txt file | Postural data obtained in the lab | Figure_1E.txt |
| Bill of Materials for Figure 2 (.xlsx) | Source of each product specified in the spreadsheet | Bill_of_materials_Figure_2.xlsx |
| Papilyawn setup. doc | Written by AF | Figure_2_Papilyawn_assembly_instructions.pdf |
| DXF (Laser cut) | Designed by AF | Back_wall_Figure_2.dxf |
| DXF (Laser cut) | Designed by AF | Bottom_Figure_2.dxf |
| DXF (Laser cut) | Designed by AF | Front_Figure_2.dxf |
| DXF (Laser cut) | Designed by AF | Shelf_Figure_2.dxf |
| DXF (Laser cut) | Designed by AF | Sides_x2_Figure_2.dxf |
| DXF (Laser cut) | Designed by AF | Top_Figure_2.dxf |
| STL (3D print) | Designed by AF | Light_ring_Figure_2.stl |
| Python script for Pico control of vibrating disks in IW experiments | Written by JF | Main.py |
| All antennograms | Output files from R (Conversion of raw DLC coordinates to 4 channels) | Figure_3_all_antennograms_D_iulia.PDF<br>Figure_3_all_antennograms_H.melpomene.PDF |
| csv | Output from DLC | Figure_2B_Dryas_iulia_D34.csv<br>Figure_2B_H.melpomene_M20.csv |
| R script | Written by AF | Figure_2B_D_iulia.R<br>Figure_2B_H.melpomene_M20.R |
| R script | Written by EM | Figure_1B.R |
| csv | Output from video tracking | Pm_Figure_1B.csv<br>Raw_Figure_1B.csv |
| R script | Written by EM | Figure_1C.R |
| csv | Output from video tracking | Raw_Figure_1C.csv |
| R script | Written by EM | Figure_3A.R |
| csv | Output from BORIS | AS_bouts_Figure_3A.csv |
| R script | Written by EM | Figure 3B |
| R script | Written by EM | Figure 3C.R |
| csv | Output from BORIS | AS_example_behaviour_Figure_3C.csv<br>AS_example_traces_Figure_3C.csv |
| R script | Written by EM | Supp_Figure_2.R |
| csv | Output form DLC | antennal_traces_Supp_Figure_2.csv |
| R script | Written by EM | Figure_3D.R |
| csv | Output from DLC | displacement_long_Figure_3D.csv |
| R script | Written by EM | Figure 4.R |
| csv | Output from BORIS | all_boris_Figure_4.csv<br>bout_behav_Figure_4.csv<br>indiv_arousal_prop_Figure_4.csv<br>indiv_prop_Figure_4.csv |
| R script | Written by EM | Figure_2C.R |
| csv | Output from DLC | Figure_2C_example_windows.csv<br>Figure_2C_overview_traces.csv |
| R script | Written by EM | Figure_2D.R |
| csv | Output from DLC | Figure_2D_example_windows.csv<br>Figure_2D_zoom_traces.csv |
| R script | Written by EM | Figure_2E.R |
| csv | Summary table derived from DLC antennal coordinates | Figure_2E_D34_fft_summary.csv<br>Figure_2E_D34_psd.csv |
| R script | Written by EM | Figure_2F.R |
| csv | Summary table derived from DLC antennal coordinates | Figure_2F_D34_acf.csv<br>Figure_2F_D34_fft_summary.csv |
| R script | Written by EM | Figure_2G.R |
| csv | Summary table derived from DLC antennal coordinates | Figure_2G_dry_summary_baseline.csv |
| R script | Written by EM | Figure_2H.R |
| csv | Summary table derived from DLC antennal coordinates | Figure_2H_raster_individuals.csv<br>Figure_2H_raster_windows.csv |
| R script | Written by EM | Figure_2I.R |
| csv | Summary table derived from DLC antennal coordinates | Figure_2I_hel_summary_baseline.csv |

## SUPPLEMENTARY FIGURES

**Supplementary Figure 1:**
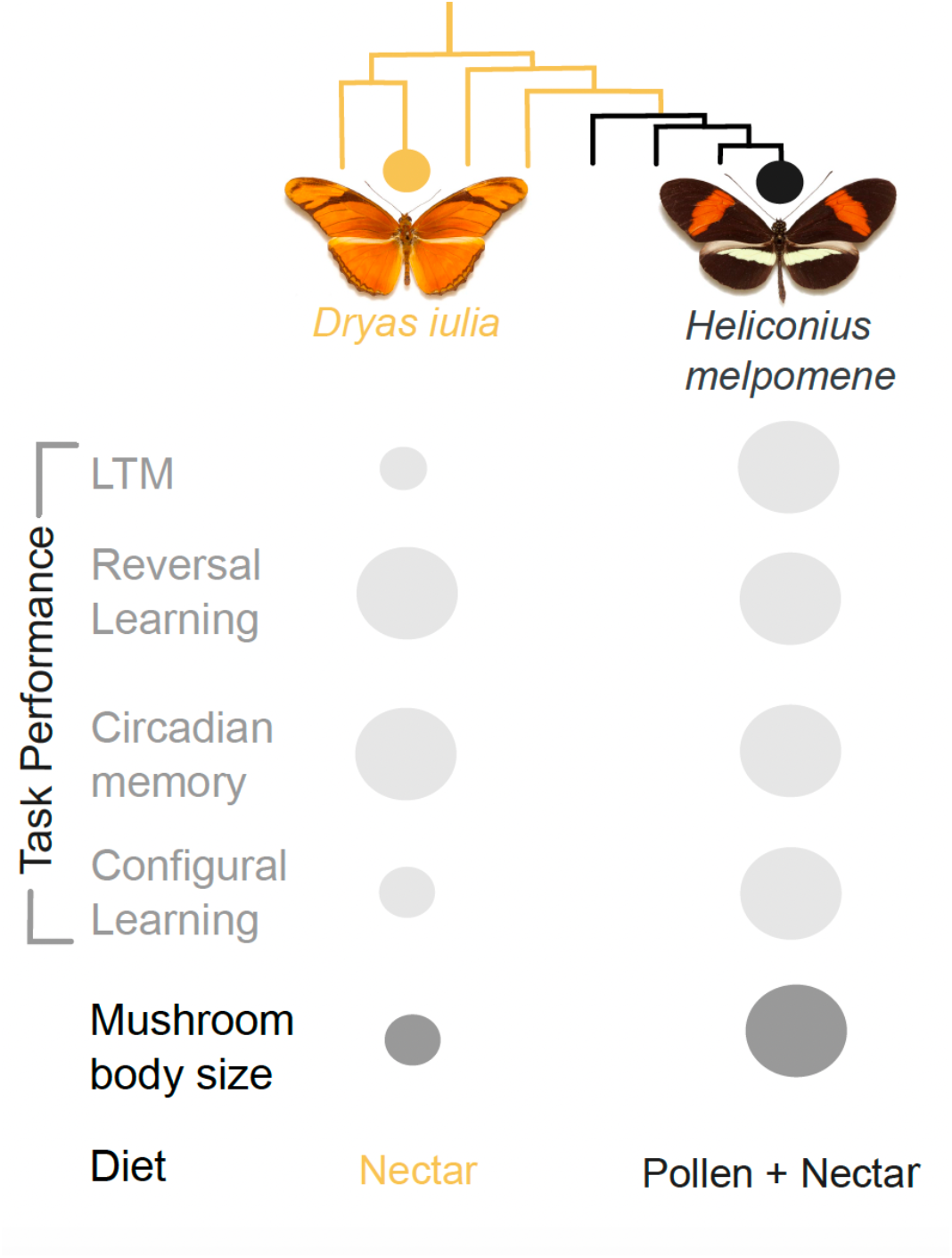
Tree showing phylogenetic relationships between *Dryas iulia* and *Heliconius melpomene*. Light grey circles represent relative proficiency in learning and memory tasks. Mushroom body size (dark grey circles) and diet are also represented.

**Supplementary Figure 2:**
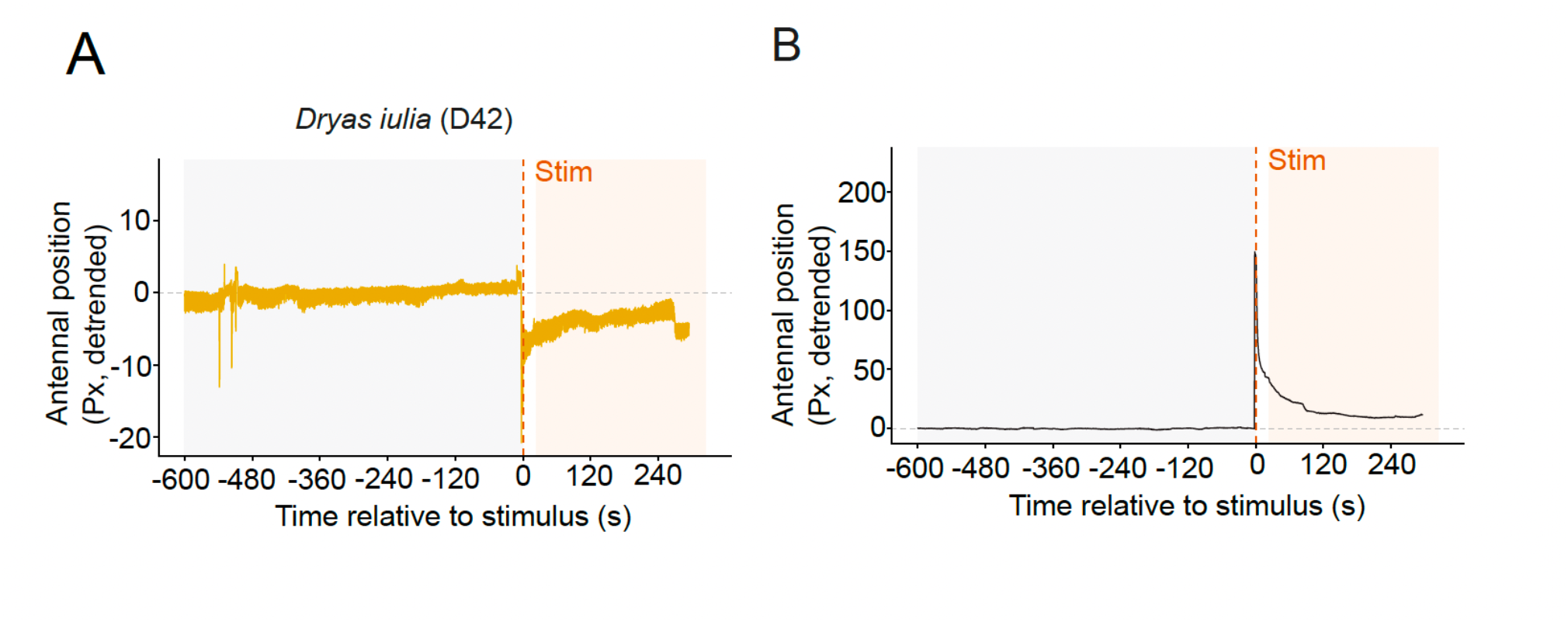
Detrended antennal positions pre- and post-vibration for representative individuals for *D. iulia* (A) and *H. melpomene* (B).

